# scBoolSeq: Linking scRNA-Seq Statistics and Boolean Dynamics

**DOI:** 10.1101/2023.10.23.563518

**Authors:** Gustavo Maganã López, Laurence Calzone, Andrei Zinovyev, Loïc Paulevé

## Abstract

Boolean networks are largely employed to model the qualitative dynamics of cell fate processes by describing the change of binary activation states of genes and transcription factors with time. Being able to bridge such qualitative states with quantitative measurements of gene expressions in cells, as scRNA-Seq, is a cornerstone for data-driven model construction and validation. On one hand, scRNA-Seq binarisation is a key step for inferring and validating Boolean models. On the other hand, the generation of synthetic scRNA-Seq data from baseline Boolean models provides an important asset to benchmark inference methods. However, linking characteristics of scRNA-Seq datasets, including dropout events, with Boolean states is a challenging task.

We present scBoolSeq, a method for the bidirectional linking of scRNA-Seq data and Boolean activation state of genes. Given a reference scRNA-Seq dataset, scBoolSeq computes statistical criteria to classify the empirical gene pseudocount distributions as either unimodal, bimodal, or zero-inflated, and fit a probabilistic model of dropouts, with gene-dependent parameters. From these learnt distributions, scBoolSeq can perform both binarisation of scRNA-Seq datasets, and generate synthetic scRNA-Seq datasets from Boolean trajectories, as issued from Boolean networks, using biased sampling and dropout simulation. We present a case study demonstrating the application of scBoolSeq’s binarisation scheme in data-driven model inference. Furthermore, we compare synthetic scRNA-Seq data generated by scBoolSeq with BoolODE from the same Boolean Network model. The comparison shows that our method better reproduces the statistics of real scRNA-Seq datasets, such as the mean-variance and mean-dropout relationships while exhibiting clearly defined trajectories in a two-dimensional projection of the data.

**Author summary:** The qualitative and logical modeling of cell dynamics has brought precious insight on gene regulatory mechanisms that drive cellular differentiation and fate decisions by predicting cellular trajectories and mutations for their control. However, the design and validation of these models is impeded by the quantitative nature of experimental measurements of cellular states. In this paper, we provide and assess a new methodology, scBoolSeq for bridging single-cell level pseudocounts of RNA transcripts with Boolean classification of gene activity levels. Our method, implemented as a Python package, enables both to *binarise* scRNA-Seq data in order to match quantitative measurements with states of logicals models, and to generate synthetic data from Boolean trajectories in order to benchmark inference methods. We show that scBoolSeq accurately captures main statistical features of scRNA-Seq data, including measurement dropouts, improving significantly the state of the art. Overall, scBoolSeq brings a statistically-grounded method for enabling the inference and validation of qualitative models from scRNA-Seq data.

## Introduction

Unveiling the mechanisms that regulate cellular decisions is a central task in systems biology. For instance, numerous efforts have been conducted to elucidate the core mechanisms that control differentiation and cell fate decision processes such as osteogenesis [1–3], haematopoiesis [4–7], dopaminergic neuron differentiation [8], early retinal development [9], and various cancer types [10–13].

The advent of single-cell RNA sequencing (scRNA-Seq) technologies has greatly enhanced the resolution with which these dynamic phenomena can be studied. As a preliminary step, most studies first determine cell identities via either clustering and subsequent manual annotation or via the direct classification of cells [14]. Furthermore, trajectory reconstruction methods [15–17] allow visualising and hypothesising how gradual changes in gene expression eventually lead to commitment to specific lineages and phenotypes. A tremendous challenge is then to identify regulatory mechanisms that control the identified dynamics of expression patterns and ultimately phenotypes.

Boolean networks are widely employed to model cellular differentiation [18–21] and fate decision [22, 23]. In these models, the activity of biological entities is represented as either active or inactive. This coarse-grained view of gene expression levels helps counter the varying levels of technical noise caused by sequencing technologies. The binary representation allows reasoning on the causal relationships between entities without having to estimate kinetic parameters or regulation thresholds, while ensuring consistency with underlying quantitative models [24]. Boolean models can predict trajectories and conclude on the impossibility of certain behaviours, optionally subject to mutations, and can encompass thousands of genes. They revealed to be a powerful and relevant modelling approach to predict combinations of genetic perturbations to control cell fate decision [25, 26].

Nevertheless, linking qualitative gene activation states with their quantitative measurements, such as count of RNA transcripts, is a delicate task with high stakes for Boolean modelling. We present scBoolSeq, which, given a reference dataset, provides a bidirectional link between scRNA-Seq and Boolean activation states.

The binary coarse-graining of scRNA-Seq, we refer to as *binarisation*, consists in assigning a qualitative active or inactive state to a gene, from one single-cell or a pool of single-cell measurements. The pools of cells usually correspond to phenotypes and other important cellular states. As Boolean models aim at predicting stability and trajectories between such cellular states, binarised data are crucial to assess their fitness with trajectories and steady states. One can easily note that the binary classification may be irrelevant in some cases, e.g., when in intermediate activation levels, or because of lacking statistical support. Therefore, it is important that binarisation methods actually result in three possible outcomes of the gene state: activate, inactive, or undetermined. However, numerous methods fully binarise transcriptome data with no regard for uncertainty or intermediate expression and the diversity of empirical pseudocount distributions [27]. RefBool [28] provided an important effort for quantifying statistical uncertainty for the binarisation and allowing intermediate states. Their approach aims at exploiting a user-defined gene expression library which serves as a proxy to take into consideration the context of the global gene expression landscape when coarse-graining data. Unfortunately this approach is only available for bulk RNA-Seq data.

The inverse operation of binarisation consists in generating RNA pseudocounts from Boolean activation states. Coupled with simulations of Boolean models, this enables generating synthetic datasets from Boolean models subject to ranges of combinations of perturbations, simulating gene knock-out or constitutive activation, for instance. Resulting synthetic scRNA-Seq data can then serve as a basis to evaluate inference methods, such as gene regulatory networks inference, trajectory inference, and Boolean model inference.

Generating single-cell and bulk RNA-Seq data has been addressed by count simulators [29–31]. With different underlying assumptions, count simulators reproduce the statistical characteristics of real datasets via parametric and semi-parametric approaches. They are capable of simulating a wide variety of scenarios and even batch effects, but generally fail at integrating information from GRN known a priori. Efforts have been made to integrate knowledge about GRNs into count simulators [32]. However, this method requires the GRN to be a directed acyclic graph, which might not be the case in general. Alternative methods rely on translating Boolean networks into non-linear Ordinary Differential Equations (ODEs). A first work in this line was odefy which presented a canonical way of transforming Boolean into continuous models [33]. More recently, boolODE was presented in the context of GRN inference method benchmarking [34, 35], introducing the addition of noise terms to make the ODEs stochastic. By building on top of Boolean networks, these approaches enable to capture the logical and dynamical relationships among the regulators. boolODE uses Hill functions to reflect the modulation of gene expression [36–38]. However, this approach relies on a considerable amount of parameters such as mRNA transcription and degradation rates, Hill thresholds and coefficients, signalling timescales, and interaction strengths. Determining these parameters is an important bottleneck as they can hardly be estimated from experimental scRNA-Seq data and need therefore to be set arbitrarily or randomly sampled. Moreover, these ODE-based generators fail to produce data with statistical properties comparable to those of real scRNA-Seq datasets.

We believe it is crucial that generated count data resemble as much as possible scRNA-Seq data to obtain fair inference benchmarks, which implies mimicking dropouts and other statistical features. scBoolSeq relies on the learning of gene-wise RNA pseudocount statistics from a reference dataset. This learning is performed in three steps: (i) the classification of empirical gene pseudocount distributions; (ii) the use of Gaussian Mixtures with up to two components as a parametric model; and (iii) the simulation of dropout events with probabilities that are inversely proportional to the expression value. scBoolSeq requires the reference dataset to be constituted of only highly variable genes (HVGs). Functions to perform this filtering are available on major scRNA-Seq analysis distributions such as Stream [15] and Scanpy [17]. By selecting HVGs after quality control, normalisation, and batch correction, one ensures that scBoolSeq’s reference reflects the underlying biological variation rather than technical noise. In addition to HGVs which are automatically selected by the designated functions in scRNA-Seq analysis environments, differentially expressed genes (DEGs) and known markers can also be incorporated to scBoolSeq’s reference in order to have a fuller image of the transcriptional landscape of the dynamic phenomenon of interest.

Thus, from the preprocessed reference dataset, scBoolSeq is able to perform two distinct complementary operations: the binarisation of a scRNA-Seq dataset with respect to the reference dataset, and the generation of synthetic scRNA-Seq from Boolean activation states, as illustrated by Fig. 1.

**Fig 1.**
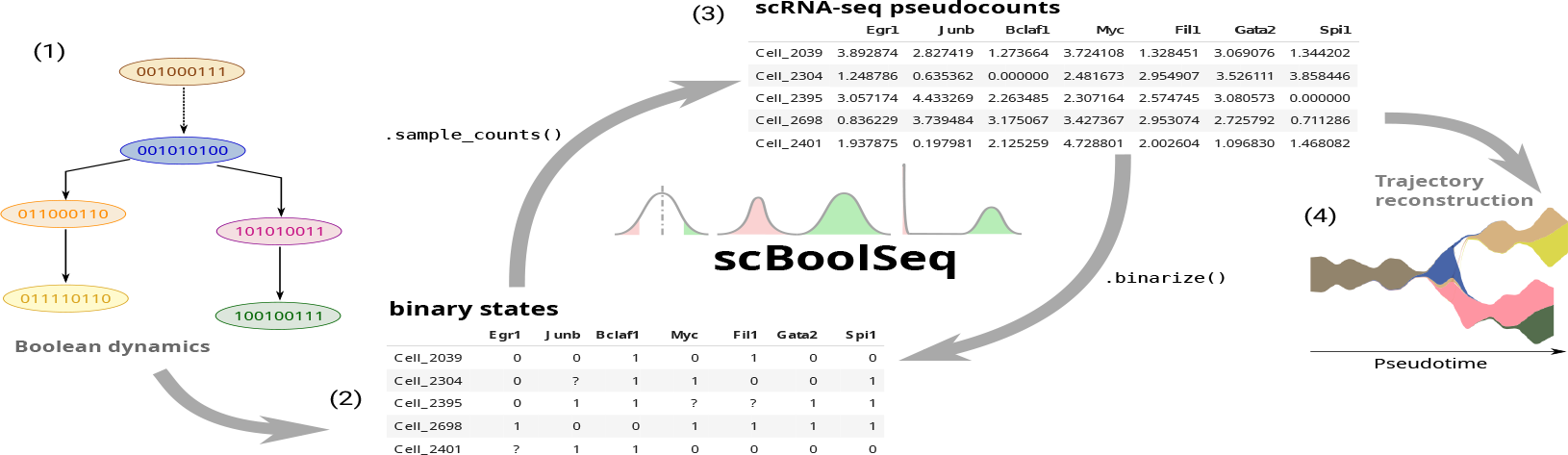
From left to right: (1) A branching trajectory constructed by merging two Boolean simulations, each one leading to a different stable state. (2) A binarised expression matrix, having genes as columns and samples as rows. (3) A pseudocount matrix (same format as the Boolean matrix). (4) A STREAM-plot reconstructing the branching trajectory from synthetic data generated from the Boolean traces [15]. scBoolSeq can be used to go from gene expression matrices (such as 3) to Boolean matrices and vice-versa.

We first show that our 3-distribution model of scRNA-Seq counts and dropouts is able to accurately reproduce the statistical characteristics of a range of scRNA-Seq datasets. For the binarisation of scRNA-Seq data, we first apply our method to a publicly available scRNA-Seq dataset of early retinogenesis. We show that scBoolSeq correctly identifies the different cell types described in the original study, defined by a minimal set of marker genes. These identities can subsequently be used in order to label cell groups found by the louvain clustering algorithm [39]. Going beyond cell type identification, we use the Boolean gene activity values determined by scBoolSeq in order to prune a mouse regulon database [40]. The resulting GRN is validated via Gene Set Enrichment Analysis performed using METASCAPE [41] which yielded numerous relevant Gene Ontology terms related to the kept genes.

Finally, we show that scBoolSeq’s synthetic scRNA-Seq data generated from Boolean traces produces both discernible trajectories when applying dimensionality reduction techniques and statistics that comparable to those of real datasets.

Overall, scBoolSeq provides an efficient method to learn statistics of a scRNA-Seq dataset and derive binarisation and synthetic generation procedures with few parameters. scBoolSeq has been implemented as an open source Python package available at github.com/bnediction/scBoolSeq.

## Results

In the following, we assume that scRNA-Seq data is preprocessed as log pseudocounts 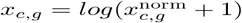, where *c* and *g* refer to cells and genes, respectively. Any size-factor based normalisation can be used, as long as it is of the form 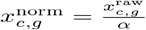 where *α* is a constant. For instance *α*_*c*_ = Σ_*g*_*x*_*c,g*_ would represent the standard library size normalisation, yielding counts/reads per million (CPM/RPM). Our methodology is applicable to alternative normalisations such as TPM (transcripts per kilobase per million reads) or RPKM (reads per kilobase per million reads). The log transformation is necessary in order to ensure the validity of the underlying parametric distributions.

### Classification of Pseudocount Distributions and Dropout Model

scBoolSeq builds on the ideas presented in [42] which seek to capture the different expression patterns across bulk RNA-Seq samples of cancer patients. By computing a series of statistical criteria, they proposed to classify empirical pseudocount distributions as bimodal, zero-inflated, or unimodal. This choice of distributions reflects the underlying hypotheses of gene activity: bimodal genes exhibit two distinct expression patters for the absence and presence of their corresponding encoded proteins. For unimodal genes, we suppose that only cells lying at tails of the distribution can be confidently inferred to be active or inactive. It also appeared that several genes show a high proportion of zeros, which are then classified as zero-inflated. Their classification method employs statistics such as mean, median, variance, dropout rate, amplitude, dip test’s p-value [43], kurtosis, density peak, and Bimodality Index [44]. In a first step, genes which do not exhibit a high enough variability or have excessive dropout rates are filtered out. Then, bimodal patterns are searched within kept genes, using a combination of statistics. Afterwards, genes with no bimodal patterns are tested for zero-inflation by looking at the empirical distributions’ density peaks. Remaining genes are classified as unimodal.

With scBoolSeq, we generalized and improved this approach to account for the specificities of scRNA-Seq data, notably their potential high dropout in gene counts, and to enable the sampling of count for reconstructed distributions in order to generate synthetic scRNA-Seq datasets from Boolean activation states. As we illustrate in S2 Fig, when applied to scRNA-Seq, the PROFILE classification algorithms show two shortcomings: (1) for genes classified as bimodal and unimodal, the dropout tends to artificially decrease their mean and inflate their variance, impeding a good characterisation of their empirical pseudocount distributions via Gaussian or two-component Gaussian Mixtures; (2) for zero-inflated genes, the classification does not result in a parametric distribution, which complicates sampling. We improved the algorithm by computing the statistics on non-zero data and propose a novel probabilistic model for dropouts in order to capture the proportion of zeros. By modelling the probability of a dropout occurring as a function of the expression level with gene-dependent parameters, we were able to reproduce the per-gene dropout rates of different reference datasets. Furthermore, we observed that, when sampling from the aforementioned parametric distributions and applying our dropout model, the zero-inflation character of certain genes as well as the excess kurtosis and skewness of unimodal and bimodal genes were globally recovered (S3 Fig).

### Probabilistic Simulation of Dropout Events

Dropouts arise from both biological (lack of transcription at measurement time) [45] and technical causes (sampling and amplification bias) [46]. For this reason, we built a probabilistic model aiming to: (i) reproduce the distribution of dropout rates across genes in the studied reference datasets; (ii) have a minimal set of gene-dependent parameters; and (iii) have a physical interpretation that accounts for the biological and technical causes of dropouts. Dropout parameters are estimated on a gene-dependent basis because empirical sampling rates exhibit gene-specific bias rather than being uniform random samples of mRNA molecules present in the cell [47]. By modelling this gene-dependent biases and simulating dropout events after sampling from parametric distributions, our dropout method mimics the physical phenomena that give rise to dropout events and generates data that reproduces the statistics of scRNA-Seq data, as illustrated by Fig. 2.

**Fig 2.**
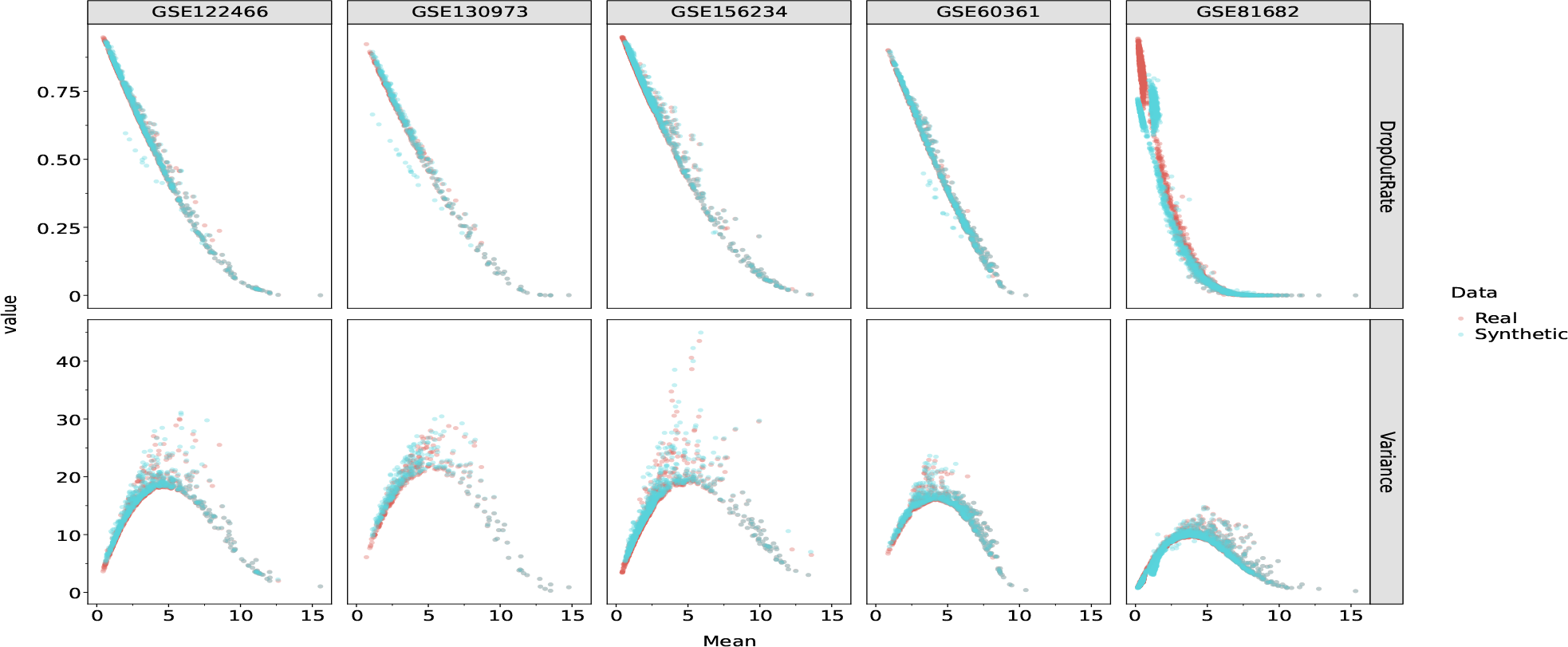
Mean - Variance, and Mean - Dropout Rate relationships of HVGs in different datasets. Each blue dot represents the average of 100 samples for a given gene.

### Dropout model

Under the hypothesis that the probability of not observing counts for a certain gene within any given cell is inversely proportional to its relative abundance, the relationship is defined as an exponential decay which has been shown to describe the mean-dropout relationship in several scRNA-Seq datasets [48]. We denote by *x*_*c,g*_ the prior pseudocount of gene *g* in cell *c* and by 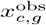 the measured pseudocount. The mathematical formulation of the proposed dropout model is of the following form:

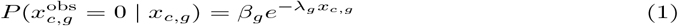

When simulating dropout events based on these probabilities, the number of dropout events for a given gene across all cells follows a Poisson-binomial distribution [49], that is the discrete probability distribution of a series of independent Bernoulli trials whose success (dropout) probabilities are not necessarily identical. This reflects our hypotheses on dropouts: for any given gene, having a dropout event for cell *i* is independent of the dropout in cell *j*, and two cells having comparable relative transcript abundances of any given gene will have similar probabilities of this gene being observed or dropped-out.

### Rate parameter

The rate parameter *λ*_*g*_ determines the shape of the exponential and thus how rapidly the dropout probabilities decay with the expression value. This parameter is learnt from the reference dataset, independently for each gene, in order to reflect the aforementioned gene-dependent sampling bias. It is calculated by setting the half-life of equation 1 to the gene’s empirical non-zero mean as follows, for each gene *g* of the reference dataset:

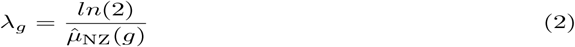

where 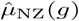 is the mean of non-zero pseudocounts of gene *g* in the reference dataset.

### Normalisation constant

The normalisation constant *β*_*g*_ is computed from sampled prior pseudocounts as the optimum value minimising the quadratic deviation between the expected dropout rate of the synthetic sample E [*τ*_*g*_] and the reference dropout rate for that gene 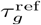 (proportion of zero entries in the reference dataset):

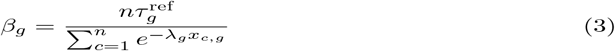

where *n* is the number of sampled cells.

This optimum is derived analytically from the expected value of a Poisson-binomial distribution. This ensures that for the same underlying non-zero distribution the dropout rate will, on average, be close to that of the reference.

S1 Fig shows an example of the distribution of rate parameters and the obtained dropout probabilities over the range of expression of a typical log-normalised scRNA-Seq dataset. Overall, we observe a trend depending on the gene pseudocount distribution category: for the same sampled value, zero-inflated genes have the highest probability of dropout, followed by bimodal genes. Genes presenting a unimodal distribution have the lowest dropout rates (and highest non-zero means) and therefore will be seldom dropped-out.

### Validation

We validated our model by sampling from the learnt parametric distributions and simulating dropouts with our exponential model of Eq. (1). We found that our method reproduces extensive statistics of these datasets, specially the gene mean-variance and mean-dropout relationships which characterise scRNA-Seq data (Fig. 2). Furthermore, the correlation profile between all combinations of mean, variance, skewness, and excess kurtosis is globally recovered (S3 Fig). We find that these correlations are only recovered when applying our dropout simulation method.

### Binarisation of scRNA-seq data

The coarse-graining scheme of scBoolSeq is based on the classification of pseudocount distribution from a reference dataset, as illustrated by Fig. 3. For each gene, cells whose expression level is high (respectively low) enough to classify it as True/active (resp. False/inactive) will be binarised whilst cells whose expression level is ambiguous will be left as undefined. As shown in Fig. 3, the category-dependent binarisation strategy causes each distribution type to have different proportions of False, True, and undetermined values.

**Fig 3.**
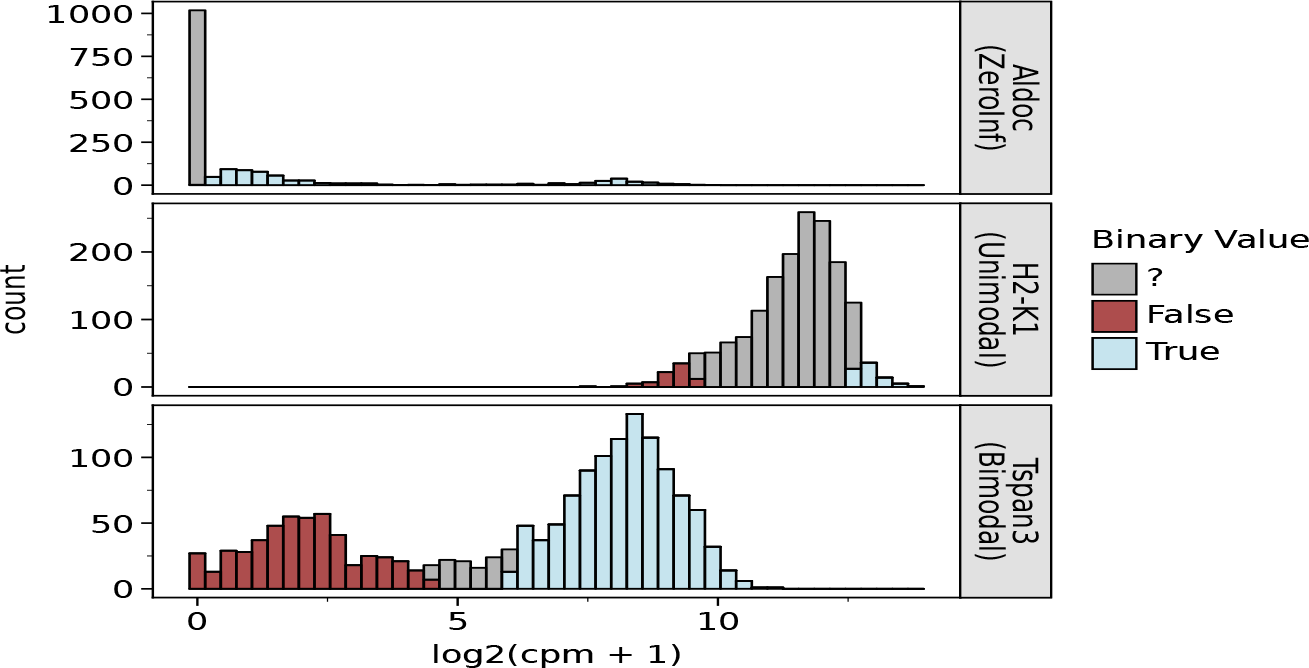
Illustration of the category-dependent binarisation allows accounting for different shapes in empirical pseudocount distributions. For each category, plots show the empirical distribution for a selected gene in the GSE81682 dataset, and the part of the values which are binarised with parameters *z* = “?” for zero-inflated case, *q* = 0.05 and *α* = 0 for unimodal and *θ* = 0.95 for bimodal.

Bimodal genes are binarised using their corresponding univariate two-component Gaussian Mixture Model (GMM), whose parameters are estimated on the reference dataset. The GMM’s density is given by Eq. 4. The model has two components denoted *C*_*i*_ which are characterised by their parameters 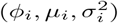. In the following, it always holds that *µ*_2_ *> µ*_1_, for every bimodal gene. Therefore, we have two components which represent cells whose transcript level can be classified as active *C*_2_ or inactive *C*_1_.

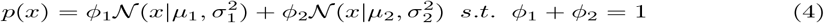

The probabilities of observation *x* belonging to each one of the two components are first calculated as detailed in Eq. (5):

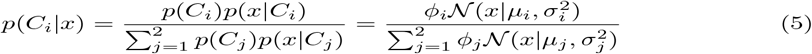

Then, the binary classification is performed according to a given confidence threshold *θ*, with 0.5 *< θ ≤* 1:

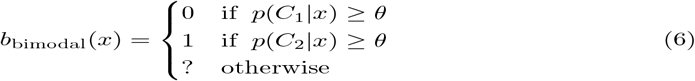

For genes classified as unimodal, we use symmetric thresholds based on two parameters: a *margin* quantile *q* (0.05 by default) and a multiplier *α* for the interquartile range IQR. These thresholds are based on Tukey’s fences for outlier detection [50], with modified defaults to binarise a small fraction of observations. Note that in Eq. (7), *Q*(*q*) represents the *q*-th quantile of the gene’s empirical distribution.

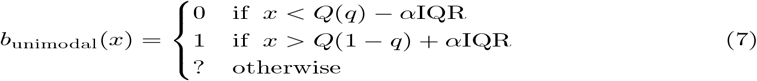

Finally, genes whose empirical pseudocount distribution is classified as zero-inflated use a zero-or-not binarisation scheme [45]. Genes having non-zero counts are classified as True whilst zero entries are classified as undetermined (parameter *z* = “?^*”*^) to reflect the uncertainty regarding the technical/biological causes of this zero, or as False (parameter *z* = 0) if considered as a signal, as suggested by [45].

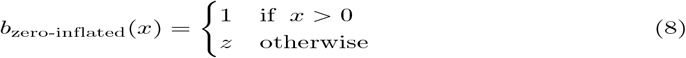

The proportion of observations classified as 0 or 1 can be approximated by Eq. (9)

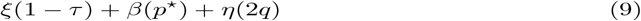

with the average proportions of binarised observations for each category normalized by the proportion of genes classified as zero-inflated, bimodal, and unimodal, denoted by *ξ, β*, and *η*, respectively, and where *τ* represents the average empirical dropout rate.

Fig. 4 gives statistics on the fraction of observations that are binarised across the selected evaluation datasets. In general, zero-inflated genes with a high dropout rate will only have a few observations binarised to 1 and most cells will be classified as undefined. Bimodal genes are binarised across most cells because the underlying Gaussian Mixture correctly describes the bimodal genes’ empirical distributions. Finally, unimodal genes will have twice the margin quantile *q* fraction of observations binarised in the case of *α* = 0 in Eq. (7).

**Fig 4.**
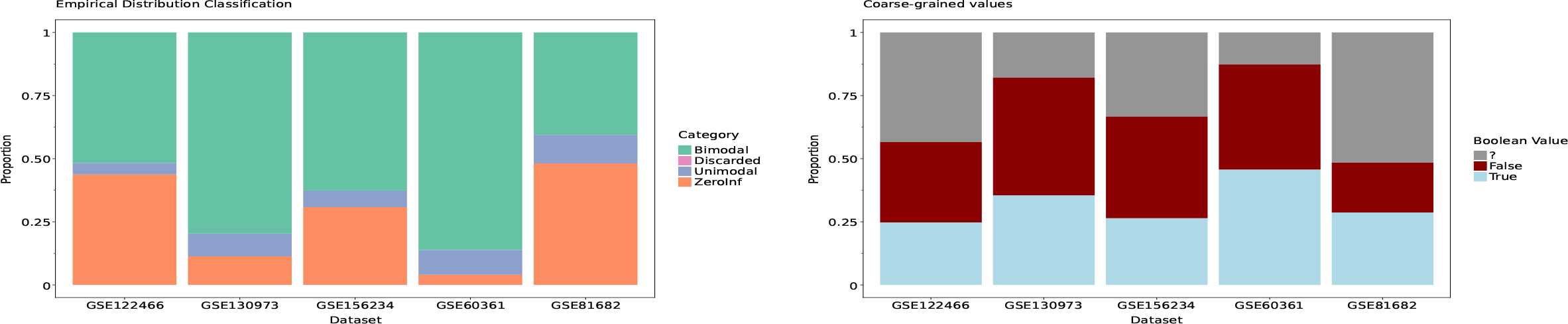
Left: Distribution of categories among the studied datasets. Right: Proportion of binarised values across datasets using the default parameters for each distribution type. These proportions are both determined by the categories and the specified thresholds. These were obtained using parameters *z* = ? for zero-inflated case, *q* = 0.05 and *α* = 0 for unimodal, and *θ* = 0.95 for bimodal. The dropout rate threshold for marking a gene as discarded was set to 0.99.

### Case study of binarisation: Early-born Retinal Neurons

We applied scBoolSeq to a publicly available scRNA-Seq dataset in order to binarise expression data and obtain a qualitative description of phenotypes. We show that the obtained qualitative profiles can serve as a basis to perform inference of Boolean networks, which can mimic the differentiation process and identify key genes and interaction involved in the dynamics.

The dataset originates from [9] (*GEO accession* GSE122466) which analysed how the diversity of cell types found in the early retina (from embryonic days 10 to 17) arises from a pool of progenitor cells. These neurons are retinal ganglion cells (RGCs), cone photoreceptors (cones), horizontal cells (HC) and amacrine cells (AC). The analyses extended previously known marker genes and showed how these appear to be organised in transcriptional waves of co-expression. Extending the original results with a mechanistic model could help formulate hypotheses regarding the underlying regulatory mechanisms of early retinogenesis. Here, we illustrate how to combine the statistical analysis of scBoolSeq to coarse-grain the expression data with prior knowledge data on transcription factor regulations publicly available in the mouse regulon database DoRothea [40] in order to build logical models which reproduce the differentiation process. Our objective is to first evaluate how the binarisation preserves the cell type classification, and how the resulting qualitative description of phenotypes enables to identify core regulations that explain the Boolean differentiation process.

### Discriminating cellular types using prior-knowledge markers

The reference study [9] considered prior knowledge markers for the cellular types at different stages of differentiation. We classified each cell according to its binarised expression profile and the markers it contains. Then, for each cellular type, we computed how many cells have the matching marker, and among them, how many match only with that cellular type. As shown in Table 1, the majority of cells per group were unambiguously identified, except for Horizontal Cells. Notice that Horizontal Cells share one marker *Prox1* with Amacrine Cells. It should be noted, that in this case, a quarter of cells have been classified using their binarisation (S4 Fig). Moreover, our classification of cells based on their binarised pseudocounts and prior-knowledge markers enables to label Louvain clusters of scRNA-Seq data, which turned out to be consistent with labels obtained using differential expression analysis by [9] (S5 Fig).

**Table 1.** A list of all the cellular types of interest, as well as the Boolean markers (cells with those genes binarised to 1/True/active) used to detect cells matching belonging to them. *N. Unambiguous Cells* represents cells that exclusively expressed the given set of markers.

| Cell Type | N. Unambiguous Cells | Perc. Total | Markers |
| --- | --- | --- | --- |
| RPC (Retinal Progenitor Cells) | 249 | 98.03% | Sox2, Fos, Hes1 |
| NB1 (Neuroblasts, first group) | 23 | 85.19% | Top2a, Prc1, Sstr2, Penk, Btg2 |
| NB2 (Neuroblasts, second group) | 27 | 81.82% | Neurod4, Pax6, Pcdh17 |
| RGC (Retinal Ganglion Cells) | 191 | 94.55% | Isl1, Pou4f2, Pou6f2, Elavl4 |
| AC (Amacrine Cells) | 81 | 67.50% | Onecut2, Prox1 |
| HC (Horizontal Cells) | 3 | 10.71% | Onecut1, Prox1 |
| Cones (Photoreceptors) | 8 | 100% | Otx2, Crx, Thrb, Rbp4 |

### Data-driven inference of Boolean models

The binarisation of scBoolSeq enables to specify Boolean dynamical properties that reflect the observed differentiation process: existence of trajectories linking (partially) binarised cellular states, including branches from pluripotent states to distinct differentiated states, as well as stability properties. Then, inference methods such as BoNesis [51, 52] can derive Boolean networks that reproduce the specified dynamics. The logical rules are derived from prior knowledge Gene Regulatory Networks (GRNs), typically extracted from TF-TF (transcription factor - transcription factor) interaction databases, possibly completed with statistical network inference from scRNA-Seq data. By employing combinatorial optimization method, BoNesis enables accessing to the sparsest models, i.e., requiring as few as possible genes to reproduce the desired trajectories and stable states. Using clustering and trajectory reconstruction methods, we applied scBoolSeq to determine a partial binary profile of 6 cellular types, namely `RPC` (progenitor), intermediate neuroblast types `NB1` and `NB2`, and final `Cones, RGC and AC` types. Note that due to the low number of cells classified as HC and their apparent distance between each others, we omitted this cellular state. The dynamical specification consisted in the existence of a trajectory from the `RPC` state to `NB1` and then to `NB2`. From the `NB2` state, three different trajectories must exist towards each of the final stable states. Moreover, we extracted from the DoRothea database a core `TF-TF` regulatory network together with target genes which have been binarised. Focusing on the largest weakly connected component, it gave a GRN with 644 genes. Then, using BoNesis, we reconstructed Boolean networks that, using the input GRN interactions, are able to reproduce the desired trajectories and stable states. See Methods section, S7 Fig, and S1 Code for details. Because the binary profiles are partials, numerous genes have no imposed binary state in several cellular states. Using BoNesis, we identified models which relies the little as possible on the dynamics of those genes with undetermined states. It resulted in pruning the input GRN to 184 genes which suffice to explain the observed differentiation process. As shown in Fig. 5(Right), gene ontology enrichment analysis, performed using Metascape [41], shows many relevant ontology terms were found among the top hits, such as mechanisms associated with pluripotency, negative regulation of cell differentiation, regulation of mitotic cell cycle, gland development, regulation of developmental growth, and embryonic organ development. Obtained models can then serve as inputs for a more thorough systems biology analysis of the biological problem.

**Fig 5.**
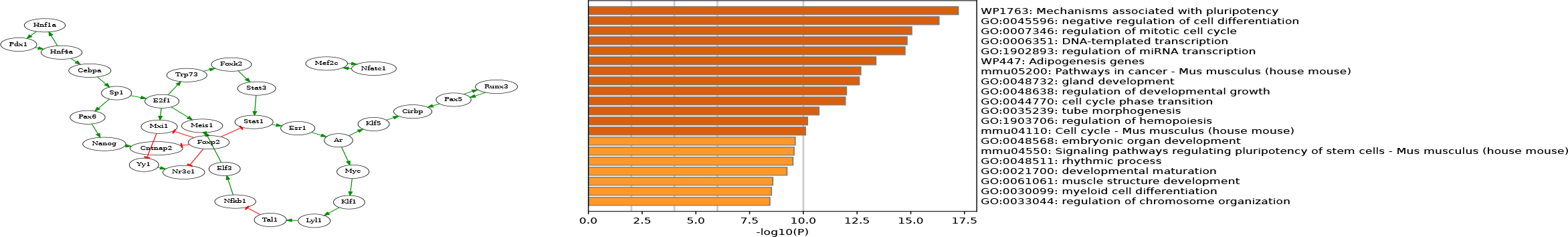
**Left**: Simplified view of the set of minimal TF-TF interactions employed in the Boolean models reproducing the differentiation process. For display, all leaf nodes with an in-degree of 1 where recursively removed from the GRN. The full filtered GRN obtained with BoNesis is provided in S6 Fig. **Right**: Top Gene Ontology Terms related to the 184 genes of the filtered GRN.

### Synthetic scRNA-seq generation biased by Boolean states

As the inverse operation of binarisation, the parametric distributions and dropout model learned per genes from a reference dataset also enable generating synthetic pseudocounts corresponding to Boolean activation states. The main principle is to perform first biased sampling from distributions whose parameters are learnt on non-zero entries of the reference dataset. In a second step, dropout events are simulated according to the gene-dependant model of Eq. (1).

Biased sampling ensures that cells in which a gene is active will exhibit higher expression (pseudocounts) than those in which it is inactive. In the case the gene follows a unimodal distribution of median *µ* and variance *σ*, the pseudocount are sampled from the half-normal distribution corresponding to the activation state (*ℋ 𝒩* (*µ, σ*^2^) for active, and *µ -* (*ℋ 𝒩* (0, *σ*^2^) for inactive). In the case of bimodal distribution, composed of two normal distributions of median *µ*_1_ *< µ*_2_ and variance *σ*_1_ and *σ*_2_, respectively, the sampling is performed from the mode corresponding to the activation state (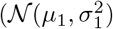 for inactive and 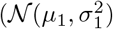 for active). Finally, in the case of zero-inflated genes, the learning from non-zero entries ensures falling back to one of the two aforementioned cases, and the dropout model learnt should reflect the inflation of zeros. The last step simulates dropouts in such a way that synthetic log-pseudocounts produced from the Boolean states will have gene-wise statistical properties closely resembling those of real scRNA-Seq data. The dropout event simulation can follow the dropout model of Eq. (1) learnt per gene, or follow an arbitrary given distribution.

### Application to artificial Boolean models

The above steps enable generating synthetic scRNA-Seq datasets from collections of binary states of genes, as it would be typically generated from the simulation of Boolean networks [53, 54]. This generation can then serve as a basis for benchmarking inference methods, by creating synthetic datasets from fixed dynamical models and evaluate the ability of inference methods to recover main features of the ground-truth model. This could notably be applied from artificial Boolean models of different scale and topology. In that cases, however, node are not directly referring to the genes of an experimental scRNA-Seq reference dataset, and one need criteria to associate a reference gene to them.

A possible approach, proposed in scBoolSeq, is to analyse the shape of the node-wise distribution of Boolean values and assign genes having similar shape. Intuitively, a gene is for instance active in most cells, it can be classified as Unimodal. Subsequently, genes which vary considerably can be considered to be Bimodal. Genes which are ubiquitously inactive with a couple exceptions (e.g., it is active in only one state of the Boolean trace) would then be zero-inflated. scBoolSeq uses scaled versions of the first four moments to classify Boolean gene distributions as unimodal, bimodal, or zero-inflated. The scaled moments of Boolean distributions are fed to a k-nearest-neighbours classifier that was trained on the scaled moments of reference dataset, using their corresponding distribution types. Afterwards a by-category bijective matching is performed in order to ensure that the synthetic scRNA-Seq distributions correctly represent the underlying Boolean dynamics.

We applied this principle on three artificial Boolean models, exhibiting different type of emerging dynamics. For each one of the models, Boolean trajectories representing the dynamics of the network were obtained as described in the next paragraphs. Afterwards multiple observations (corresponding to single cells) were sampled using scBoolSeq with a selected reference dataset (GSE81682). Then, we applied classical scRNA-Seq dimensionality reduction methods to visualise the corresponding pseudocount trajectories. Further details regarding the sampling procedure and projections can be found in the supplementary materials.

The first artificial model is a star-like network (Fig. 6a) in which a single Transcription Factor (TF) up-regulates the expression of a set of genes. This model was simulated by performing one random walk with the fully asynchronous update mode starting from the state where the node `tf` is active and all genes are inactive. The resulting trajectory is a sequence of Boolean vectors where genes progressively activate, in a random order. This gradual activation can be clearly distinguished in Figure 6b, where cells with few active genes are coloured in dark blue and cells with all genes active are coloured in light green.

**Fig 6.**
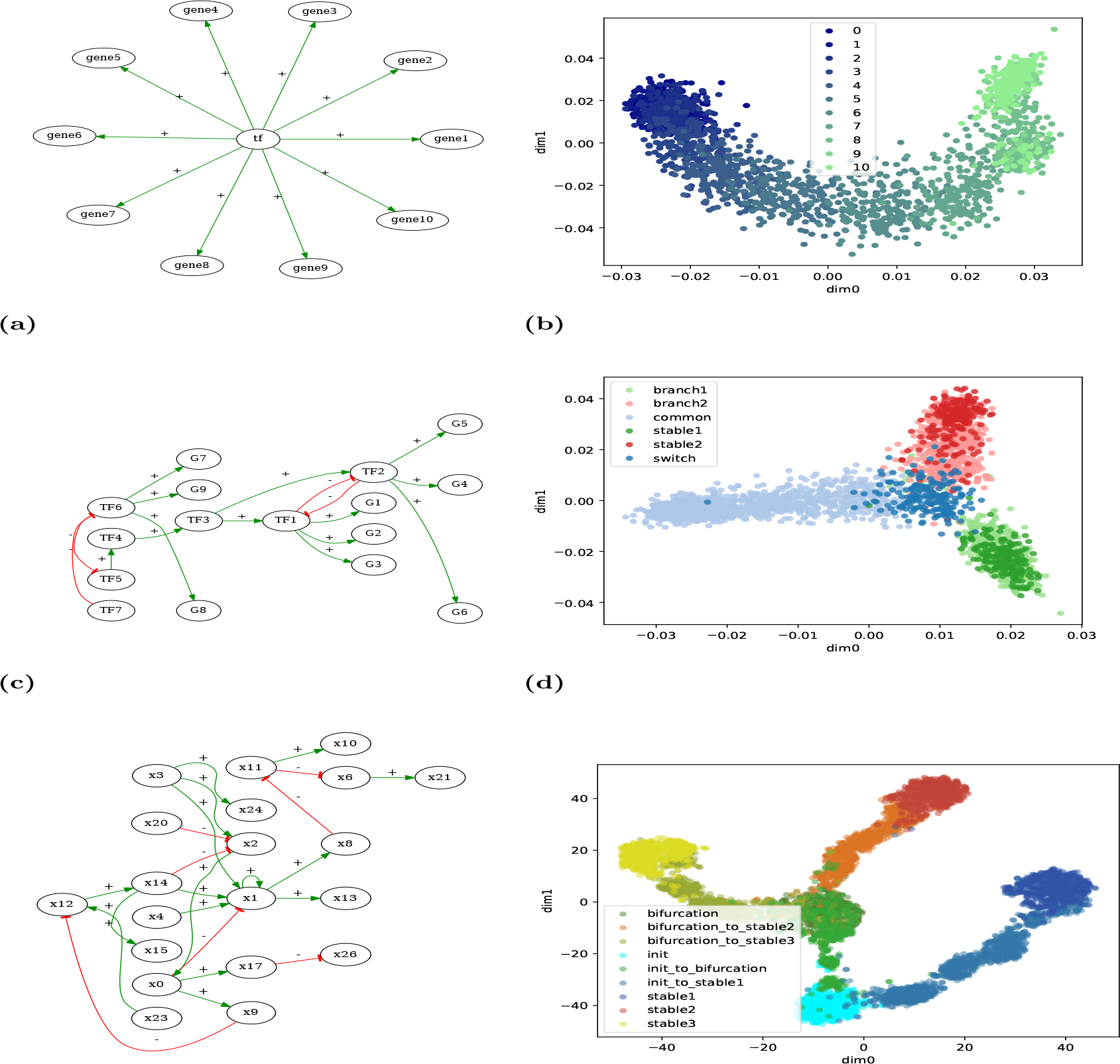
Artificial Boolean models and generated synthetic scRNA-Seq data. **Left**: Influence graphs of the Boolean models. See supplementary material for Boolean functions. **Right**: Two-dimensional projection of the synthetic scRNA-Seq data generated by applying scBoolSeq to Boolean trajectories simulated from the models on the left. Dots are labelled with a description of the Boolean state they have been generated from: for (b) it is the number of active genes; for (d) and (f) they refer to the dynamical nature of the states in the 3-branches differentiation process.

The second manually-designed model is a bistable switch which represent a simplified *cellular reprogramming* scenario (Fig. 6c) in which the cell finds itself in a steady state (light blue, labelled *common*) characterised by the activation of *TF6* which activates a small set of genes and inhibits a mutually exclusive switch. The activation of *TF7* node represents a perturbation which inhibits *TF6*, pushing the cell out of its initial state and triggering a differentiation process. At the end, one of two different stable states is reached. The third model (Fig. 6e) is a three-stable switch which has been designed automatically from random scale-free topology and such that it exhibits a two-level differentiation process: from an initial state three stable states are reachable, with an intermediate branching state giving access to two of them. In both cases, we generated Boolean trajectories covering the differentiation branches from the initial states. These trajectories remain apparent in the projections of generated scRNA-Seq data (Fig. 6d and Fig. 6f).

### Comparison with BoolODE

Given an artificial Boolean network, the tool boolODE [34] is capable of producing synthetic pseudocount datasets which exhibit clearly defined trajectories when applying dimensionality reduction techniques such as *SNE*. However, the generated dataset do not exhibit observed statistics of experimental scRNA-Seq dataset.

Fig. 7 provides comparisons between datasets generated by boolODE and scBoolSeq from one of the largest curated model of the benchmark of [34], a Boolean network of human gonadal sex determination (GSD) [55]. It has two main fixed point attractors of biological interest, namely Sertoli cells and granulosa cells which correspond to male and female phenotypes. We notably compared the mean-variance and mean-dropout profiles of generated data with different dropout models, as proposed by both tools. Besides the dropout rate being constant, the mean-variance relationship of boolODE appears to be at very different scale than typical scRNA-Seq data (Fig. 2). It should be noted that when enforcing a constant dropout rate with scBoolSeq, the resulting dropout-mean profile is not constant as 0 values can still be sampled from learnt pseudocount distributions: gaussian distribution can give non-zero probabilities to negative values, which are corrected as 0. This is not the case with boolODE because of the noise added to ODE-simulated values, which prevents generating values being exactly 0.

**Fig 7.**
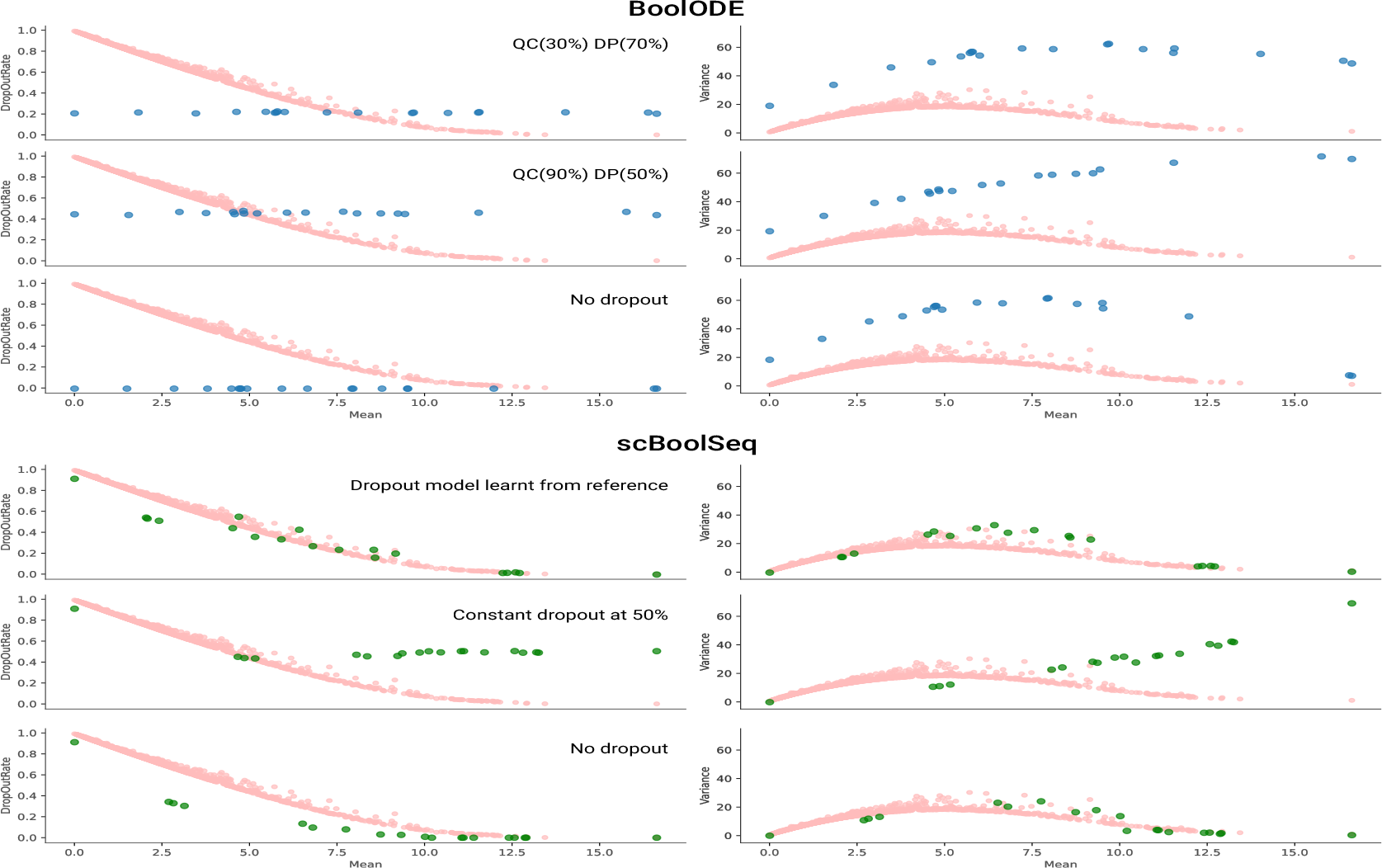
Comparison of the per-gene Mean-Variance and Mean-DropOutRate profiles of reference dataset GSE122466 (red), boolODE (blue), and scBoolSeq (green). *QC* represents the quantile below which boolODE simulates dropouts with a constant probability *DP*.

### Implementation and usage

scBoolSeq has been implemented in Python on top of `pandas` [56], `statsmodels` [57], and `scikit-learn` [58] libraries. Fig. 8 shows basic usage of scBoolSeq to perform binarisation and synthetic data generation. Future engineering work will focus on leveraging the `AnnData` [59] Python package for handling large datasets that cannot be fit in RAM. Furthermore, using `AnnData` within scBoolSeq will allow its integration in the `scverse` [60] computational ecosystem for single-cell omics data analysis.

**Fig 8.**
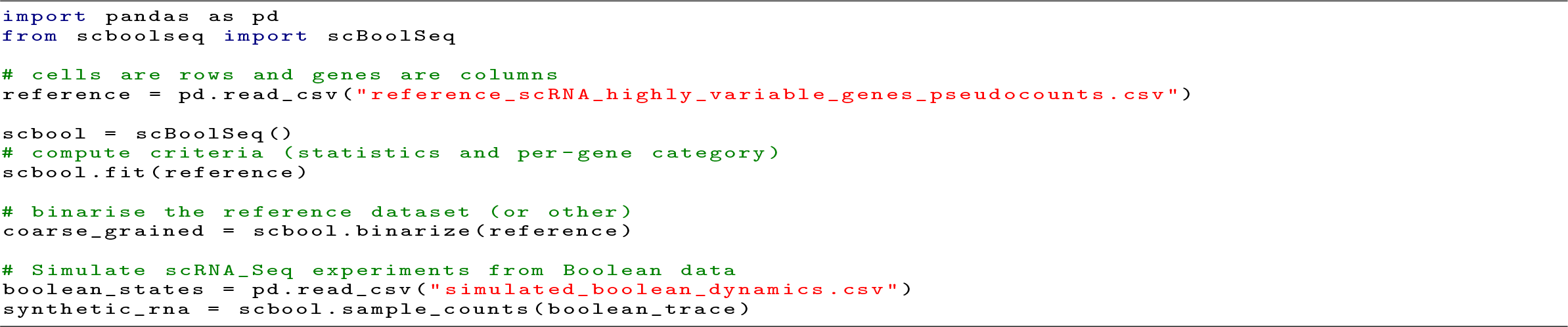
Python code snippet showing basic usage of scBoolSeq for binarisation and synthetic data generation from reference scRNA-Seq data and Boolean states

scBoolSeq is distributed as a standard Python package, and is integrated in the CoLoMoTo Docker distribution [61], which facilitates the accessibility of tools related to Boolean and logical models, and the reproducibility of related computational analyses.

## Discussion

We introduced scBoolSeq, a novel method which provides a bidirectional link between scRNA-Seq data and Boolean Models. Our method builds on the classification of gene empirical pseudocount distributions into unimodal and bimodal distributions proposed by [42], that we extended with a probabilistic gene-dependent dropout model. We showed that the resulting characterization suffices to capture the main statistical features of real scRNA-Seq data. Then, scBoolSeq offers both the ability to binarise scRNA-Seq datasets and the ability to generate synthetic pseudocounts from binary states of genes.

### From pseudocounts to binary states

We illustrated on a concrete application how the binarisation offered by scBoolSeq can be employed to process scRNA-Seq data in view of performing inference of Boolean networks, which are logical models of gene activity dynamics. First, scBoolSeq coarse-graining method allows identifying cellular types of interest by detecting the presence (i.e. activation) of known marker genes. In addition to this, combining the binarised gene activity with community detection techniques could help to find previously unknown marker genes (genes which are binarised as active only in certain clusters and are not found in the literature). Then, coupled with a prior GRN, the deduced set of Boolean functions constitute a set of hypotheses that can guide future wetlab experiments in order to unveil the core regulatory mechanism of early retinogenesis.

It should be stressed that the binarisation of scBoolSeq can result in undetermined state when there is not enough statistical evidence for a binary classification. We believe that the fact that not all genes (and cells) cannot be classified with binarisation is good sign that the method enables discriminating cells in extreme state from cell in transient state, for which a fully binary view may not be adequate.

One should note however that determining the activity of a gene based on its transcript level is a strong hypothesis. Methods such as VIPER [62] aim at adding information about each protein’s regulon to better infer protein activity. Moreover, chromatin accessibility and other epigenetics information can also help to refine the binary classification.

### From binary states to pseudocounts

Another major contribution of scBoolSeq is its method for generating synthetic scRNA-Seq data from Boolean gene activation states by biased sampling from learned pseudocount distributions on a reference dataset. We showed that scBoolSeq provides a significant improvement over boolODE as it produces synthetic scRNA-Seq data whose statistical characteristics (mean-variance and mean-dropout profiles) closely resemble those of real data. In addition to this, scBoolSeq allows simulating any arbitrary distribution of gene-wise dropout rates. This represents an unprecedented contribution as it allows measuring the sensitivity of inference methods to the dropout rate distributions of scRNA-Seq datasets.

By offering the capability to generate synthetic scRNA-Seq datasets from ground-truth Boolean models with realistic statistical features, we believe that scBoolSeq is a clear asset for that generating benchmarks for the evaluation of various inference methods, such as GRN inference, trajectory reconstruction, and data-driven Boolean network inference.

## Methods

### Boolean networks and dynamics

A *Boolean network* on nodes {1, …, *n*} is a function *f* : 𝔹^*n*^*→* 𝔹^*n*^ mapping binary vectors of dimension *n* to themselves, where 𝔹 = {0, 1} is the Boolean domain. For each node *i ∈* {1, …, *n*}, we write *f*_*i*_ : 𝔹^*n*^*→* 𝔹 the *i*-th component of *f*, which is the Boolean function of node *i*. A Boolean vector **x** *∈* 𝔹^*n*^ specifies a Boolean state for each component of the network, and is called a *configuration*.

The *influence graph* of a Boolean network *f* is a directed signed graph, noted *G*(*f*), whose vertices are the nodes of the Boolean network. The influence graph captures the dependencies of Boolean functions, and corresponds to union of Jacobian matrices of *f* on configuration. Formally, there is a positive edge for node *j* to *i* (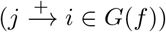) in the influence graph if and only if there exists a configuration **x** *∈* 𝔹^*n*^ such that

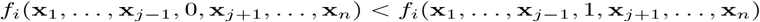

There is a negative edge for node *j* to *i* (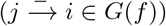) in the influence graph if and only if there exists a configuration **x** *∈* 𝔹 ^*n*^ such that

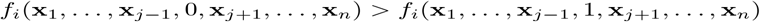

Note that it is possible to have both edges 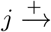 *i* and 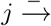 *i* in a same influence graph. If it is the case for *G*(*f*), then the Boolean network *f* is said to be non-monotone.

Otherwise, *f* is *locally monotone*.

A *trajectory* of a Boolean network *f* is a sequence of configurations **x**^1^, · ·, · **x**^*k*^ that can be computed according to a given *update mode*. For instance, the synchronous mode computes trajectories such that any two successive configurations **x**^*m*^, **x**^*m*+1^ are such that **x**^*m*+1^ = *f* (**x**^*m*^); the fully asynchronous update mode computes trajectories such that any two successive configurations **x**^*m*^, **x**^*m*+1^ differ on only one node *i*, and verify that 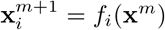. The *most permissive* update mode [24] computes all the trajectories that are binarised from any asynchronous trajectory of multivalued and quantitative model compatible with the Boolean network. In general, it allows much more trajectories than synchronous and (general) asynchronous modes, which fail to capture trajectories of different class of quantitative systems, including incoherent feed forward loops [24].

A configuration **x** *∈* 𝔹^*n*^ is a *stable state* if *f* (**x**) = **x**, i.e., it is a fixed point of *f* . A configuration **x** *∈* 𝔹^*n*^ belong to an *attractor* of *f* under a given update mode whenever for any possible trajectory from **x** to another configuration **y**, there exists a trajectory going back to **x**. Stable states are particular cases of attractors.

### Inference of Boolean networks from influence graph and dynamical properties

From an influence graph 𝒢 and a set of dynamical properties, the tool BoNesis [51, 52], available at github.com/bnediction/bonesis, allows inferring all the locally-monotone Boolean networks *f* having their influence graph enclosed by 𝒢, i.e., *G*(*f*) *⊆*, 𝒢and that posses the input dynamical properties. The dynamical properties supported by BoNesis include the existence of most permissive trajectories between partially specified configurations, and stable state properties of (partially specified) configurations. A partially specified configuration specify a Boolean state for a subset of nodes. In that case, BoNesis is free to complete the unspecified nodes with any Boolean state. BoNesis also allows specifying optimization objectives to filter solutions, notably to enumerate only sparser models, i.e., with the smallest influence graphs.

We employed BoNesis to infer Boolean networks from scRNA-Seq scBoolSeq binarisation (see next section), and to generate artificial Boolean networks which possess multi-stability and branching behaviors from randomly generated scale-free influence graph (S1 Code).

### Case Study: Early Born Retinal Neurons

We performed the analyses on the scRNA-Seq dataset of lane 1 of GSE122466. The main steps hereafter denoted in paragraphs refer to the analyses performed in their homonymous Jupyter Notebooks provided in S1 Code.

### Highly Variable Gene Selection

For this part we used the software STREAM [15]. We took the count matrix of the first replicate (Identified with the prefix `Lane_1` in their index). We performed standard quality control, with the same parameters as the analyses of the original article. Cells expressing less than 200 genes where discarded, as well as genes expressed in less than 3 cells. We selected the 1648 most highly variable genes and appended to them the two marker genes which were reported in the article but were not selected as being highly variable (*Rbp4, Pou4f2*).

### Retinal Differentiation Clustering and Metadata

In this part we took the aforementioned Highly Variable Genes (HVGs) and performed the scBoolSeq distribution learning with *θ* = 0.75 to have a higher the amount of binarised observations on bimodal genes. We then used the instance to binarise the HVGs across all cells. We then identified cells matching the markers described in the original article. About 25% of all cells where labelled in this process. Subsequently, cells matching more than one set of markers were discarded. The only pair of phenotypes which presented more than a couple ambiguous cells where Amacrine Cells (AC) and Horizontal Cells (HC) which had 23 cells matching both marker signatures. This was expected given that cellular types where defined with only two markers and one of them *Prox1* is shared. Having a larger (and preferably disjoint) set of markers could resolve this ambiguity. We used scanpy [17] to perform louvain clustering on the log pseudocount HVGs, with the number of neighbours set to 15. With this analysis, 11 distinct clusters were found. A small cluster of cells (cluster 10 in the notebooks) was discarded as it was determined to be an unknown cluster of unknown Retinal Ganglion Cell-like U/RGC. Our Boolean analysis also found this isolated cluster to express signature genes of RGCs. Finally, clusters where labelled using the majority label of cells whose Boolean identity matched the markers. Most clusters had absolute majorities (85%, 98%) except for one (Cluster 3 had 53.84% of cells voting NB2, and 34.61% voting AC: It was labelled NB2). These labels where used as metadata in order to perform trajectory inference.

### Trajectory Inference

Using STREAM we performed trajectory inference, using the aforementioned cluster labels as metadata. We obtained a well-defined trifurcating trajectory which is distinguishable on two dimensions. We set the root (starting point) to be Retinal Progenitor Cells (RPCs) and the three final points to be the Cones, Retinal Ganglion Cells (RGC), and Amacrine Cells respectively. Cells associated with these terminal nodes of the inferred graph where taken to be representative of their corresponding phenotypes. For the two groups of neuroblasts (NB1 and NB2), cells within the two quartiles *Q*(.25), *Q*(.75) of the root node’s pseudotime where chosen as representative of these transient phenotypes. This yields a total of 133 RGC, 79 NB1, 17 NB2, 109 AC, 78 RPC, and 69 Cones that were used to infer the Boolean model.

### Binarisation of scRNA-Seq data

We binarised all HVGs across all cells and employed the metadata obtained from the previous trajectory inference step to retrieve cell groups. We defined meta-observations by aggregating each group, using the mode as summary statistic. We further selected genes having non-null variance, which reduced the original 1650 genes to only 1426. We only retained binarised genes present in the mouse regulon database DoRothEA [40], that is 1263.

### Boolean Model Inference

Having our binarised observations and selected genes, we defined our GRN using DoRothEA [40]. DoRothEA gives a confidence score to each one of the interactions, based on the number of supporting evidence in different sources. In decreasing order, these levels are: *A,B,C,D,E*. We decided to exclude interactions with low supporting evidence, so we filtered out levels *D,E* and considered only levels *A,B,C*. With these filtered interactions, we extracted the core TF-TF network which we define to be the biggest strongly connected component of the departing graph. This core TF-TF network has 157 nodes. We then obtained the subgraph induced by these 157 core transcription factors and the binary genes comprising our observations. This yielded a GRN with 728 nodes. We tested and found that this GRN was not weakly connected. We extracted the biggest weakly connected component which contained 633 nodes. This weakly connected component was given to BoNesis as the domain of Boolean Networks to consider, and specified the desired trajectories and stable states using the specification given in S7 Fig.

## Supporting information

**S1. Fig.**
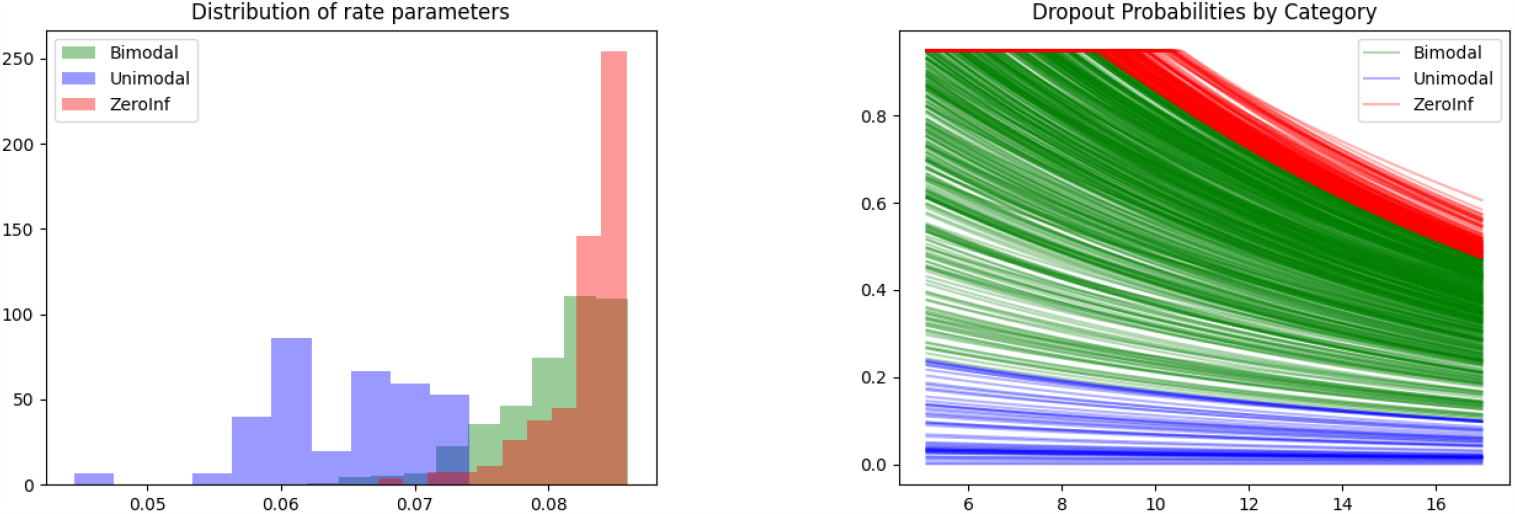
Example of distribution of rate parameters and dropout probabilities learn scBoolSeq. **Left**: Distribution of rate parameters *λ* estimated on dataset GSE122466. **Right**: Dropout probabilities computed between the minimum and maximum values of a sample from the parametric distributions corresponding to the same dataset. Each line corresponds to an individual gene.

**S2. Fig.**
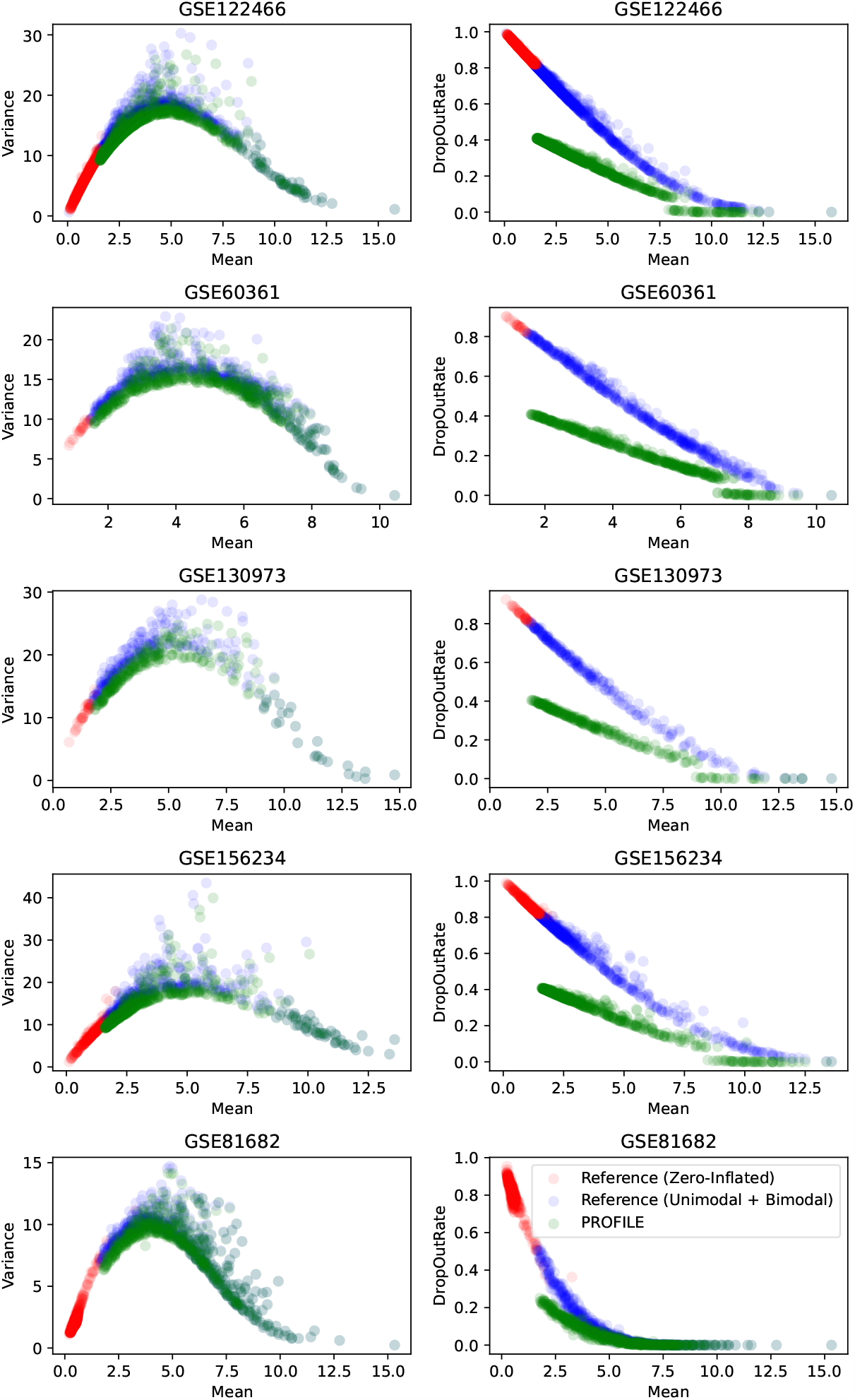
Mean - Variance and Mean - DropOutRate relationships of HVGs using PROFILE parametric distibutions for bimodal and unimodel genes on selected scRNA-Seq datasets. Each green point represents the average of 100 independent replicates with the same sample size reference dataset.

**S3. Fig.**
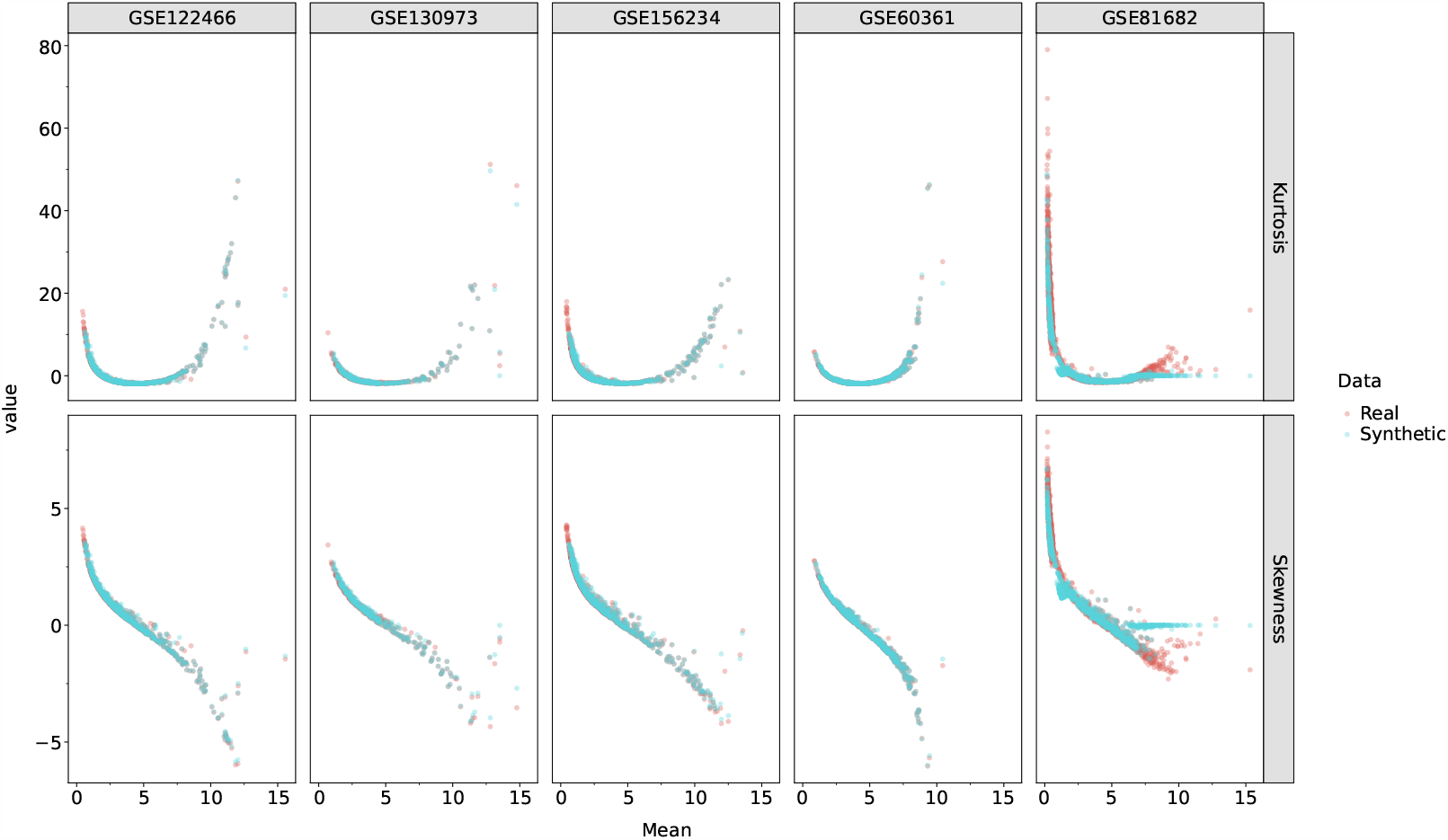
Correlation between higher moments of real pseudocount data and from data generated from distributions and dropout model learnt by scBoolSeq on selected scRNA-Seq datasets.

**S4. Fig.**
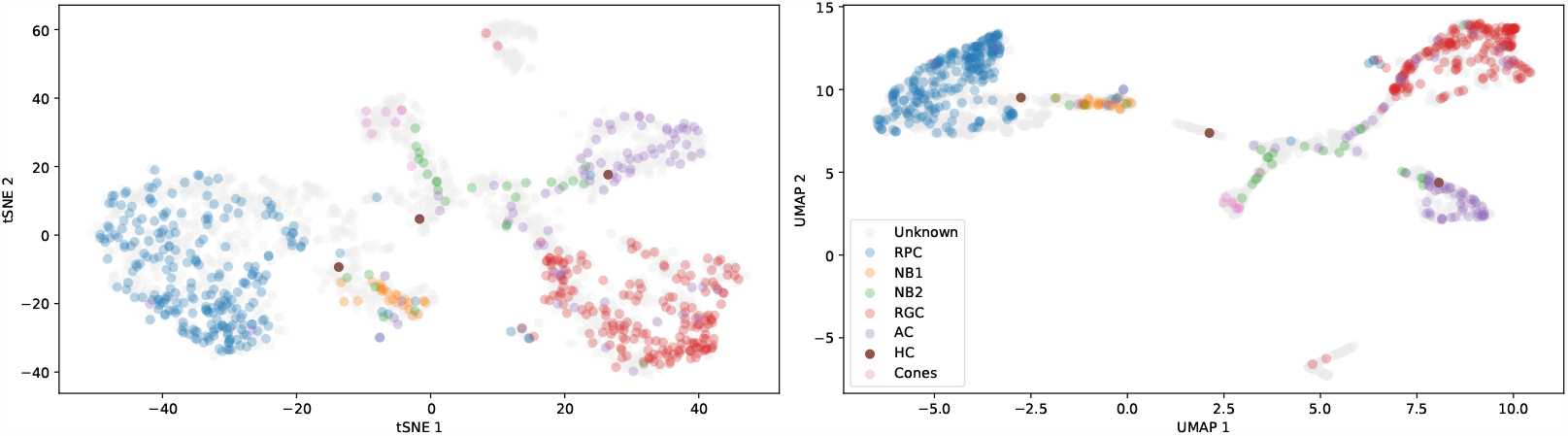
Position of cells classified using scBoolSeq binarisation and prior-knowledge markers. t-SNE and UMAP projections trained on the top 25 principal components (log pseudocount matrix). Colours indicate cell identities determined by binary value of known markers (see Table 1 of main text).

**S5. Fig.**
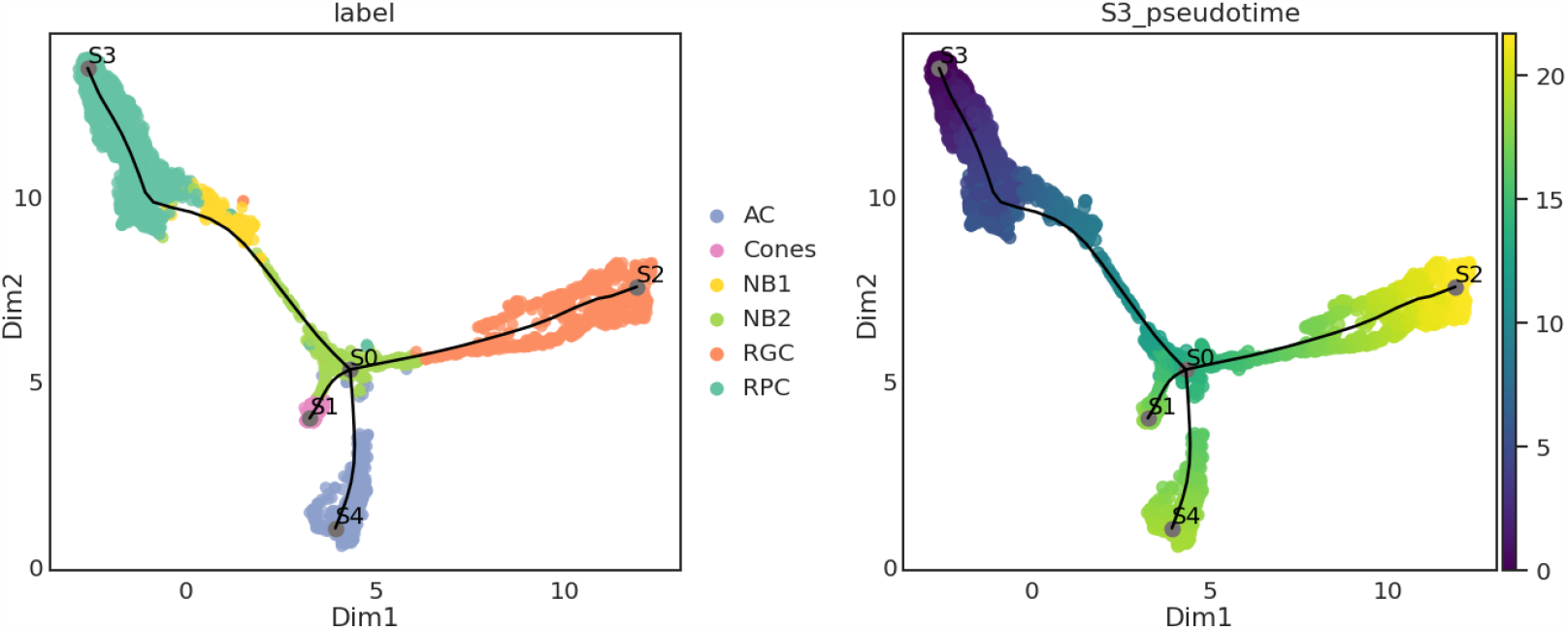
Result of trajectory reconstruction using STREAM on early-born retinal neurons scRNA-Seq data. UMAP projection of the first 25 principal components to 3 dimensions (only 2 are shown). The cluster labels are determined by the majority label of unambiguous cell types identified via scBoolSeq binarisation.

**S6. Fig.**
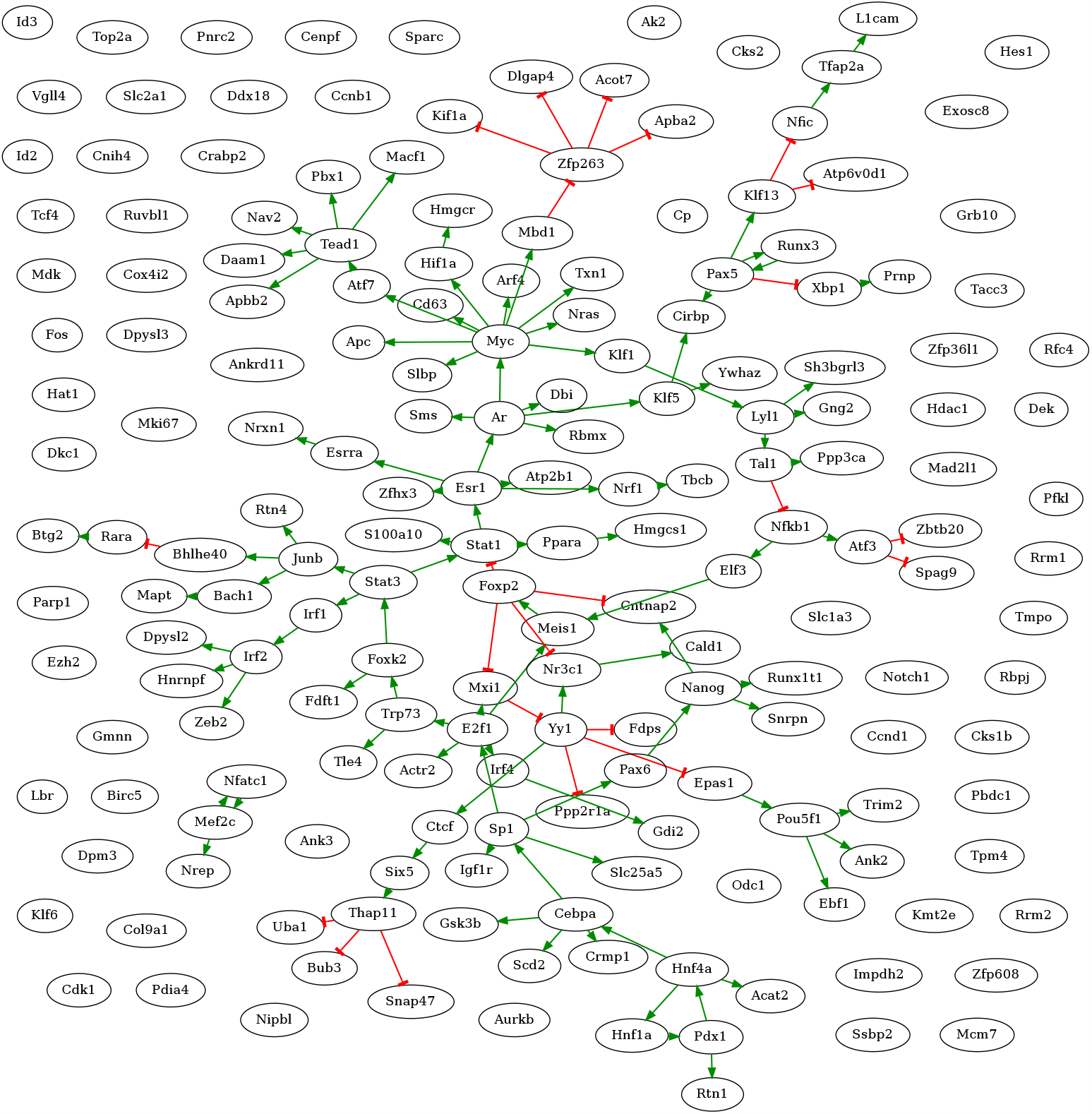
nfluence graph of sparser Boolean networks learnt using BoNesis from qualitative dynamics of case study obtained with scBoolSeq binarisation. This graph comprises 184 nodes forming Boolean networks that can reproduce the Boolean dynamics of early-born retinal neurons differentiation process. This graph is a subgraph of the input DoRothEA TF-TF interaction database. Green arrows indicate positive regulations, red arrows indicate negative regulations. Nodes without predecessors indicate nodes with constant function in the Boolean networks. Thus, the Boolean state of these nodes is identical in all stable states, and is in opposite state in the precursor state `RPC`.

**S7. Fig.**
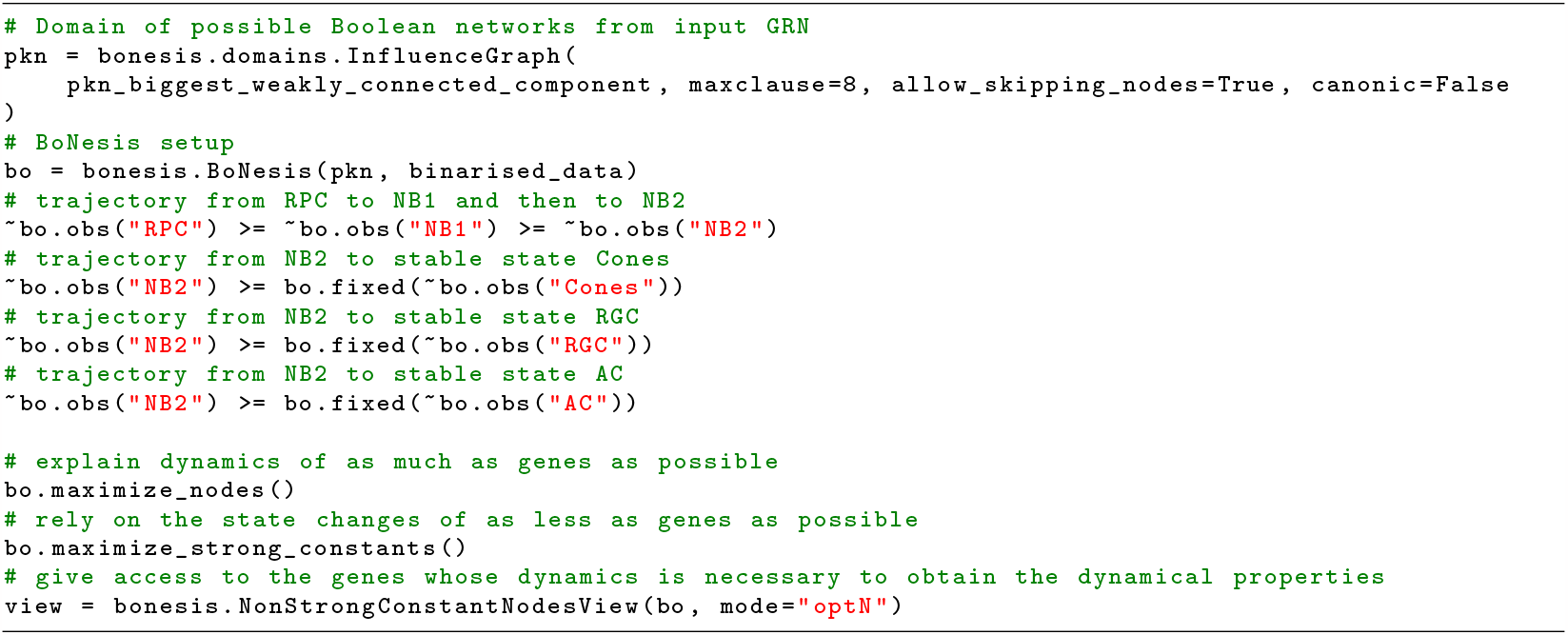
Python code snippet showing usage of BoNesis for the inference of Boolean networks for the retinal differentiation case study. See S1 Code/3. - Retinal Differentiation BN Inference for full pipeline.

**S1 Code. Code and notebooks for reproducing binarisation case study and synthetic data generation** scBoolSeq source code is available at github.com/bnediction/scBoolSeq. The Python package can be installed using conda or pip; see link for instructions. Notebooks for demonstrating scBoolSeq usage and reproducing the case studies presented in this paper can be visualised and downloaded at nbviewer.org/github/bnediction/scBoolSeq-supplementary.

